# Sex-chromosome-dependent aging in female heterogametic methylomes

**DOI:** 10.1101/2024.07.31.605993

**Authors:** Marianthi Tangili, Joanna Sudyka, Fabricio Furni, Per J. Palsbøll, Simon Verhulst

**Affiliations:** Groningen Institute for Evolutionary Life Sciences, University of Groningen, Groningen, The Netherlands; Institute of Environmental Sciences, Jagiellonian University, Kraków, Poland; Center for Coastal Studies, Provincetown, Massachusetts, USA

**Keywords:** DNA methylation, sex-dependent aging, avian sex chromosomes, female heterogametic

## Abstract

Recent research in humans and both model and non-model animals has shown that DNA methylation (DNAm), an epigenetic modification, is one of the mechanisms underlying the aging process. DNAm-based indices predict mortality and provide valuable insights into biological aging mechanisms. Although sex-dependent differences in lifespan are ubiquitous and sex chromosomes are thought to play an important role in sex-specific aging, they have been largely ignored in epigenetic aging studies. We characterized the genome-wide distribution of age-related CpG sites from longitudinal samples in two avian species (zebra finch and jackdaw), including for the first time the avian sex chromosomes (Z and the female-specific, haploid W). In both species, we find a small fraction of the CpG sites to show age-related changes in DNAm with the majority of them being located on the haploid, female-specific W chromosome where DNAm levels predominantly decrease with age. Age-related CpG sites were overrepresented on the zebra finch but underrepresented on the jackdaw Z chromosome. Our results highlight distinct age-related changes in sex chromosome DNAm compared to the rest of the genome in two avian species, suggesting this previously understudied feature of sex chromosomes may be instrumental in sex-dependent aging. Moreover, studying the DNAm of sex chromosomes might be particularly useful in aging research, facilitating the identification of shared (sex-dependent) age-related pathways and processes between phylogenetically diverse organisms.

## Introduction

The underlying causes of aging, the decline in biological functioning over time, remain an enigma. Phenotypic and molecular biomarkers that serve as proxies of biological age (Xia et al., 2017) have been instrumental in unraveling aging mechanisms. In this context, indices based on age-dependent epigenetic marks have shown an unrivaled capacity to predict chronological and biological age (Pal & Tyler, 2016). Such indices are mostly based on age-related changes in DNA methylation (DNAm) at CpG sites; sites in the genome where a 5’ cytosine is followed by a guanine (Moore et al., 2013). Indices of chronological age based upon the degree of DNAm at certain CpG loci (‘epigenetic clocks’) have opened novel, pivotal windows into the complex landscape of age-associated phenotypes. Numerous epigenetic clocks have been developed in recent years in humans (Horvath, 2013; Levine et al., 2018; Marioni et al., 2015), as well as in model (Han et al., 2020) and non-model animals (Lu et al., 2023; Tangili et al., 2023), and even plants (Gardner et al., 2023). DNAm contributes to the regulation of gene expression (Moore et al., 2013) and is known as one of the most stable epigenetic modifications, possible due to its important roles in embryonic development, cellular identity, and genomic stability (Bird, 2002). However, age-related changes in DNAm still occur but are restricted to a relatively small subset of CpG sites across the genome (Horvath, 2013). The study of changes in the level of DNAm with age across a broad range of species holds the potential to identify fundamental sites, sequences, and processes related to aging conserved across diverse taxa. Casting the net widely will enable identifying aging pathways shared among species, or classes, that likely are fundamental to aging (Moqri et al., 2024).

Sex differences in age-related pathology and lifespan are ubiquitous across the animal kingdom, where the homogametic sex in female heterogametic species lives on average 7.1% longer (Xirocostas et al., 2020), suggesting a link between sex chromosomes and aging. Despite the link between sex and age, most epigenetic aging studies have so far routinely ignored the sex chromosomes, with a few notable exceptions only in humans (G. Li et al., 2022; Lund et al., 2020). Employing a novel, robust pipeline we identified age-related CpG (AR-CpG) sites where the degree of DNAm correlates with age among blood samples from captive zebra finches (*Taeniopygia castanotis*) and wild jackdaws (*Corvus monedula*). We employed longitudinal data, i.e. two samples collected from each individual during its lifespan, to characterize the distribution of AR-CpG sites across the genome. We included, for the first time, the avian sex chromosomes (Z and the female-specific haploid W) to assess if the observed distribution of AR-CpG sites across chromosomes deviates from null expectations. Our expectation was that if sex chromosomes play a key role in aging this would be evident in the prevalence of AR-CpG sites on the sex chromosomes.

## Materials and Methods

### Sample origin

Zebra finch blood samples were collected during a long-term experiment where birds were housed in outdoor aviaries (320 × 150 × 210 cm) each containing single sex groups of 18–24 adults (Briga et al., 2017). Twenty longitudinal samples were collected between June 2008 and December 2014 from ten known-age individuals (six males and four females) sampled twice during their lifetime at an average interval of 1,470 days between the two sampling points. Jackdaw blood samples were collected from individuals of a free-ranging population breeding in nest-boxes south of Groningen, the Netherlands (53.1708°N, 6.6064°E). We analyzed 22 longitudinal blood samples collected during 2007 – 2021 from 11 known age adults (five males and six females) with an average sampling interval of 2,429 days. For both species, the precise age of sampled individuals was known because individuals were followed since birth (Table S1). Samples were collected in EDTA buffer and stored in −80°C in glycerol buffer.

### Whole genome bisulfite sequencing

We extracted total-cell DNA using innuPREP DNA Mini Kit (Analytik Jena GmbH+Co) from 3μl (nucleated) red blood cells according to the manufacturer’s protocol. Whole genome sequencing was performed by The Hospital for Sick Children (Toronto, Canada) where paired-end Illumina next-generation sequencing (150bp) was carried out on either an Illumina HiSeqX™ (12 zebra finch samples) or an Illumina NovaSeq™ sequencer (eight zebra finch samples and 22 jackdaw samples). Libraries were prepared using the Swift Biosciences Inc. Accel NGS Methyl Seq kit (part no. 30024 and 30096) and the DNA was bisulfite converted using the EZ-96 DNA Methylation-Gold kit from Zymo Research Inc. (part no. D5005) as per the manufacturer’s protocol and libraries were subsequently PCR amplified.

### Bioinformatic processing

Sequences were trimmed using Trim Galore! v. 0.6.10 (Krueger et al., 2023) in paired-end mode. Visual quality controls of the data were carried out before and after trimming using FastQC v. 011.9 (Andrews, 2010) and MultiQC v. 11.14 (Ewels et al., 2016). Because the Swift Biosciences Inc. Accel NGS Methyl Seq kit was used for library preparation, the first ∼10 bp showed extreme biases in sequence composition and M-bias, so after checking the M-bias plots, the first 10 base pairs (bp) were further trimmed from each sequence.

Alignments were performed using Bismark v. 0.14.433 (Krueger & Andrews, 2011) using the Bowtie 2 v. 2.4.5 alignment algorithm (Langmead & Salzberg, 2012) for both in silico bisulfite conversion of the reference genomes and alignments. For zebra finch, trimmed reads were aligned against the in silico bisulfite converted zebra finch (*Taeniopygia guttata*) reference genome (GCA_003957565.4, Rhie et al., 2021) and the average mapping efficiency was 64.5% (SD: 3.22). Because the jackdaw reference genome did not contain an assembled W chromosome, the jackdaw bisulfite sequencing data was aligned to an *in silico* bisulfite converted Hawaiian crow genome (*Corvus hawaiiensis*, GCA_020740725.1, Rhie et al., 2021, the closest relative species with an assembled W chromosome) the average mapping efficiency was 64.7% (SD: 0.98).

### Functional annotation of AR-CpG sites

The GFT-formatted annotation files for the zebra finch (GCF_003957565.2) and the Hawaiian crow (GCF_020740725.1) were converted into BED12 format using the University of California, Santa Cruz (UCSC) utilities *gtfToGenePred* and *genePredToBed* (available at https://hgdownload.soe.ucsc.edu/downloads.html#utilities_downloads). AR-CpG sites were annotated using the tool *annotateWithGeneParts* from the R 4.1.2 package *genomation* 1.4.1 (Akalin et al., 2015). This tool hierarchically classifies the sites into pre-defined functional regions, i.e., promoter, exon, intron, or intergenic, hereon referred to as annotation categories. The predefined annotation categories were based on the annotation information present in the BED12 files accessed with the *genomation* tool *readTranscriptFeatures*. Annotations were performed in a hierarchical manner (promoter > exon). Subsequently, a customized R script was employed to integrate the annotation results of AR-CpG sites with their respective annotation category information.

### Gene ontology enrichment analysis

The gene ontology (GO) enrichment analysis for the genes in or near which we identified AR-CpG sites was conducted using the *ClueGo* plug-in v. 2.5.10 (Bindea et al., 2009) for *Cytoscape* v. 3.10.2 (Shannon et al., 2003 and was conditioned on including at minimum three associated genes. We applied a two-sided enrichment-depletion test employing the Benjamini-Hochberg correction for multiple testing (Benjamini & Hochberg, 1995) and GO terms were considered significantly enriched if they had a corrected *p*-value was at 0.05 or less.

### Statistical assessment

We conducted *χ^2^* tests using the R package *janitor v.*2.2.0 (Firke, 2023) to compare the observed and expected number of AR-CpG sites in each species and applied exact binomial tests using the R package *stats* (R Core Team, 2023) to compare the observed and expected number of AR-CpG for each chromosome. The expected number of AR-CpG sites for each chromosome was inferred as the product of the observed number of CpG sites on each chromosome and the species-specific frequency of AR-CpG sites for the whole genome, which was 0.008% for the zebra finch and 0.006% for the jackdaw. We applied a Benjamini-Hochberg correction to control for the false discovery rate to account for multiple hypothesis testing (Benjamini & Hochberg, 1995).

We assessed the observed and expected number of AR-CpG sites per annotation category using a *χ^2^*test. The proportion of expected AR-CpG sites per annotation category was estimated as the proportion of all CpG sites that passed the initial filters (Steps 1, 2, 3 in our pipeline, Results) located in each annotation category. For all *χ^2^* tests, we used an exact binomial test as a *post hoc* test to assess observed vs. expected occurrences. All statistical tests were conducted using the CRAN R (4.1.2).

To assess whether age-related CpG sites were non-randomly distributed across genomic regions, we performed an enrichment analysis for both species (separately) based on the genomic coverage of each region. We first calculated the total length (in base pairs) of each region type per chromosome type (autosomes, W, and Z). The observed number of AR-CpGs in each region was then compared to the expected number based on that region’s relative genomic coverage. Enrichment or depletion was tested using Fisher’s exact test by comparing observed and expected CpG counts in the focal region versus all other regions. The results were expressed as odds ratios with *p-values*, with enrichment defined as an odds ratio >1 and depletion as <1. The terms “over-represented” and “underrepresented” are used to refer to sites that are more or less numerous than expected by chance on either specific chromosomes or in annotation categories if AR-CpG sites were uniformly distributed across all CpG sites in the genome.

## Results

Here we present the steps for the development of the pipeline employed to characterize the distribution of AR-CpG sites (Fig.1a).

<u>Step 1.</u> We applied a minimum mean coverage per site of ≥20 for zebra finches and ≥25 for jackdaws. Since the optimal solution with respect to the coverage filtering has to do with the distribution of the initial coverage, these thresholds were chosen based on the distribution of the initial coverage, with median values of 6 for zebra finches and 7 for jackdaws. This step depends on the quality of sequencing data and can be tailored to the coverage distribution of a particular dataset.

<u>Step 2.</u> We then selected CpG sites with an average DNAm across all samples between 40% and 60% that are likely to have a higher absolute change in DNAm over time.

<u>Step 3.</u> We the selected only the sites that were shared by the majority of the samples to increase the site detection probability across the whole population. In our datasets, that was >=15 samples for autosomes (75% of the samples in the zebra finch, 68% in the jackdaw) and >=6 samples for the W chromosome for the zebra finch (75% of female samples) and >=8 for the jackdaw (67% of female samples), as it is present only in females in both species.

<u>Step 4.</u> For each sequenced CpG site we generated a Person’s correlation coefficient between DNAm at the focal sampling point and delta age (the difference between an individual’s chronological age at the time of sampling and its average age calculated for both samplings, van de Pol & Wright, 2009) to account for between and within subject effects and correlated the Person’s correlation values with the average sequencing coverage per CpG site over all samples. To include the sites with the highest correlations between DNAm at the focal sampling point and delta age (negative or positive as we want to include methylation increases and decreases with age across all ranges of coverage) we selected the top 5% sites for every 20 units of coverage.

Since mtDNA was not found to contain any AR-CpG sites when included in the initial analysis, it was excluded from subsequent analyses. This newly developed pipeline to identify AR-CpG sites across the genome can be applied to (i) an array of different species, (ii) on DNAm data of varying qualities (in terms of sequencing coverage) and (iii) for which a limited number of individuals sampled longitudinally is sufficient. The robustness of our pipeline was confirmed by the high correlation between the chromosome-wise number of AR-CpG sites generated with different cutoff parameters (Fig. S2 and S3). This robustness stems from numerous factors. Firstly, we analyzed whole-genome DNAm data, meaning that we had the opportunity to investigate the change in DNAm of CpG sites throughout the entirety of all of the chromosomes, including the sex chromosomes. Moreover, as previously stated, the longitudinal nature of our data makes it ideal to identify within-individual changes in DNAm with age. Another strength of our method is that we take extra steps to ensure that we not only identify CpG sites whose DNAm changes with age, but CpG sites that have extreme changes (in the same direction) in DNAm that are shared by the majority of the samples in the population. Additionally, we use the metric of DNAm correlated with Δage (the difference between an individual’s chronological age at the time of sampling and its average age calculated for both samplings) instead of chronological age, meaning that individuals with larger timespans between sampling points have a higher influence on the identification of AR-CpG sites. A larger time interval between samplings exerts more pronounced changes in DNAm patterns over time, thereby enhancing the detectability of AR-CpG sites. Lastly, we observe comparable yet not identical patterns of AR-CpG site location in the genomes of our two study species which is to be expected as they belong to the same class but are not very closely related phylogenetically. Because of all the above, we maintain confidence in the strength and robustness of our results.

In zebra finches, 0.008% of CpG sites (1,468/18,906,252) and in jackdaws 0.006% (1,218/21,835,349) showed age-related DNA methylation changes (Fig. 1).We did not detect AR-CpG sites in the mitochondrial genome, but AR-CpG sites were present on most nuclear chromosomes (Fig. 2).

**Figure 1.**
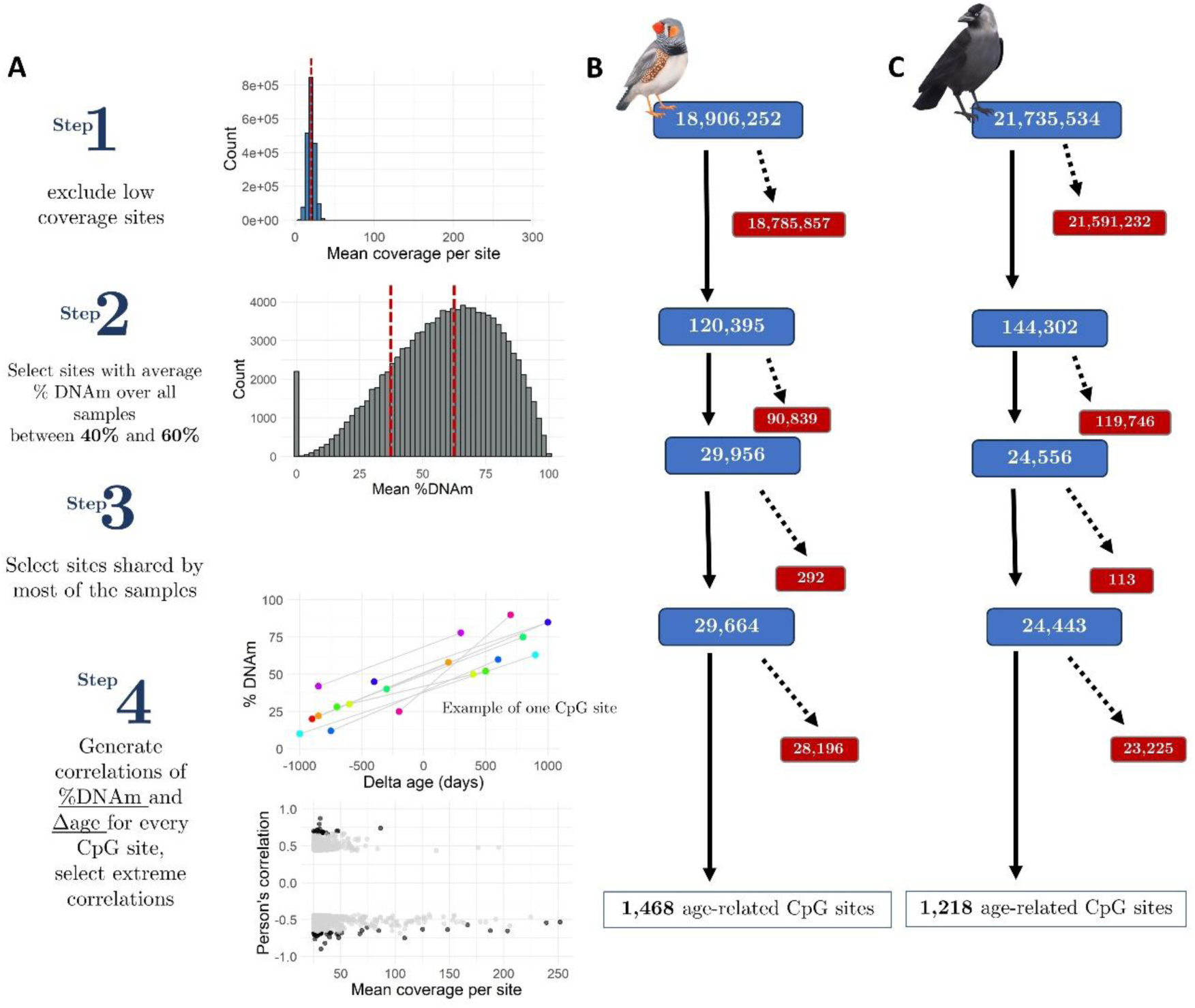
Graphical representation of the pipeline for identification of AR-CpG sites from longitudinal, whole genome bisulfite sequencing (WGBS) data for zebra finches and jackdaws (**A**). Number of CpG sites remaining (blue panels) and lost (red panels) after each step in the pipeline for the zebra finch (**B**) and jackdaw (**C**).

**Figure 2.**
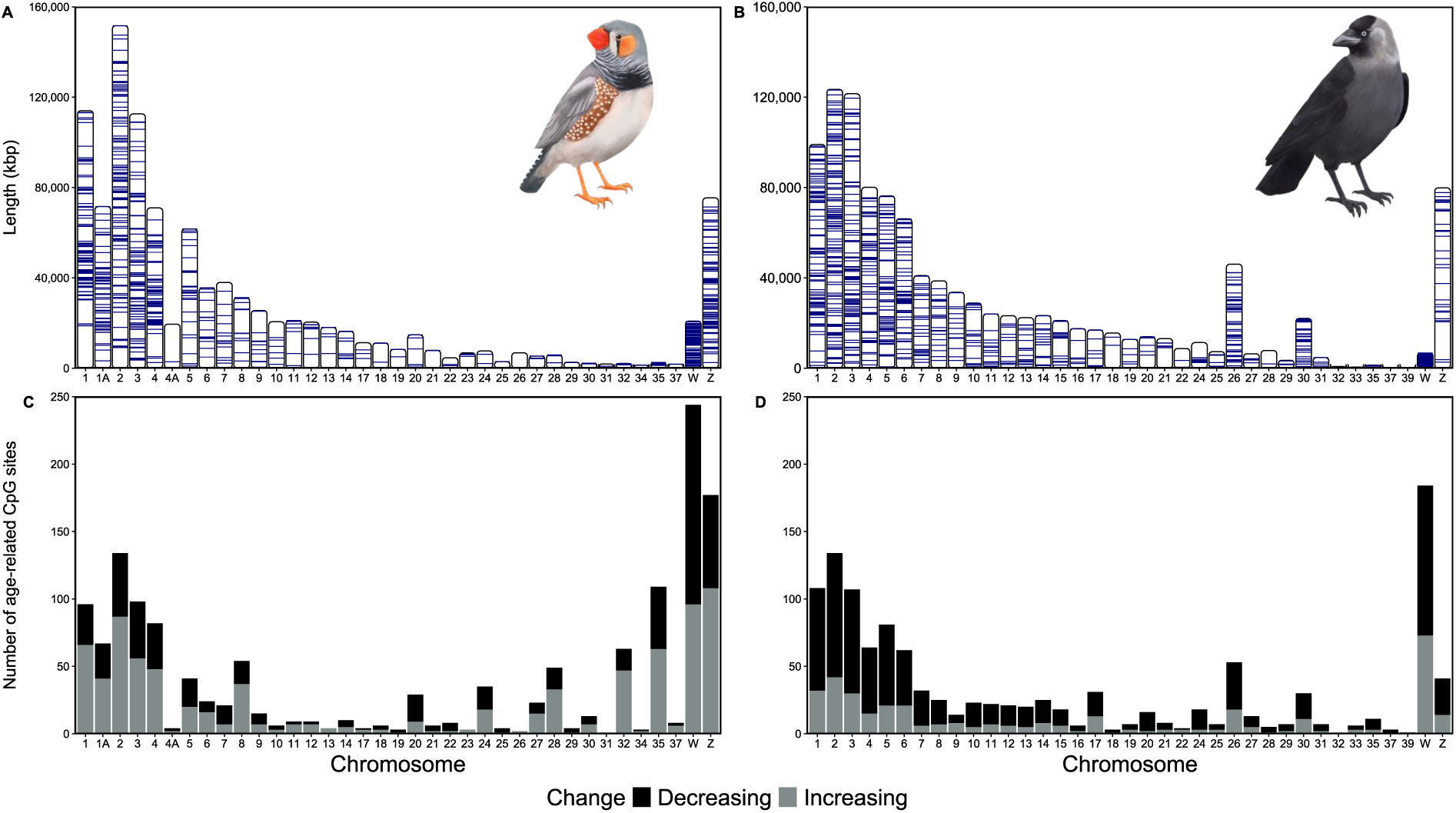
Location of AR-CpG sites across chromosomes and direction of change in DNAm. Position of each AR-CpG site per chromosome for (**A**) the zebra finch and (**B**) the jackdaw. Counts of AR-CpG sites on each chromosome whose level of DNAm increased (grey) or decreased (black) with age in the zebra finch (**C**) and the jackdaw (**D**). No AR-CpG sites were detected on zebra finch chromosomes 15, 16, 33, 36 and jackdaw chromosomes 23, 34, 36, 38. The actual AR-CpG site positions for both species can be found in (Tangili et al., 2024).

The majority of AR-CpG sites were located on the W chromosome and occurred at significantly higher frequencies than expected by chance in both species (zebra finch: Fig.3a, 17% of all AR-CpG sites, *p*<0.0001, Table S2; jackdaw: Fig.3b, 15% of all AR-CpG sites, *p*<0.0001, Table S3). The AR-CpG site density on the Z chromosome was elevated in the zebra finch (12% of all age-related sites; χ^2^= 2,622, df=36, *p*<0.001, Fig.3) and depressed in the jackdaw (3% of all AR-CpG sites; χ^2^=2,243, df=35, *p*<0.001, Fig.3). The density of AR-CpG sites also varied among the autosomal chromosomes. In the zebra finch, AR-CpG sites were overrepresented on six chromosomes and underrepresented on 22 chromosomes (Table S2). The corresponding numbers in the jackdaw were six and 10 respectively (Table S3). In the zebra finch autosomes, DNAm level increased with age on 43% of AR-CpG sites (n=632; Fig. 2C), which contrasts with the jackdaw, where DNAm increased with age on 24.5% of AR-CpG sites (n=298; Fig. 2D).

**Figure 3.**
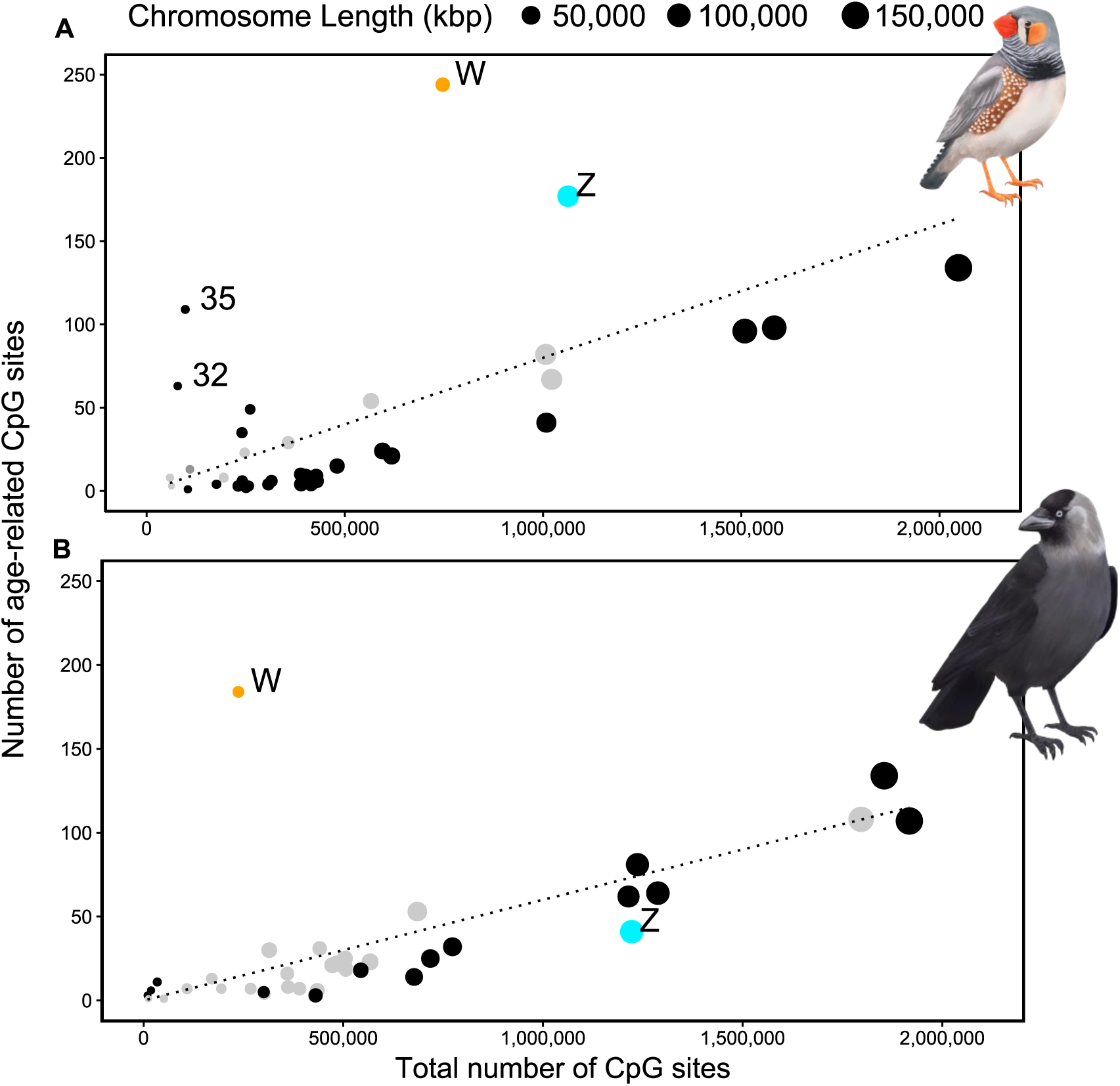
Number of AR-CpG sites in relation to total CpG sites per chromosome. (**A**) Zebra finch and (**B**) jackdaw. The diameter of each data point is proportional to chromosome length in kilobase pairs (kbp). Black data points denote autosomal chromosomes with an AR-CpG density that deviated significantly from the null expectations (dotted line).

AR-CpG sites were predominantly located in intergenic regions in both species, followed by introns, promoters, and exons in the zebra finch and introns, exons, and promoters in the jackdaw (Fig. 4A, D). The observed distribution of AR-CpGs deviated significantly from null expectations (zebra finch: χ^2^=5,655, df=3, *p*<0.001, jackdaw: χ^2^=7.98, df=3, *p*=0.04). For the zebra finch AR-CpG site distribution deviated significantly from null expectations also when considered separately for each annotation category (Table S4A), whereas in the jackdaw, only AR-CpG sites in promoters were significantly more numerous than expected (Table S4B). AR-CpG sites were absent from exons in the W chromosome in the jackdaw, while there were three AR-CpGs located in exons in the zebra finch (Fig. 4B, F, Fig. S1).

**Figure 4.**
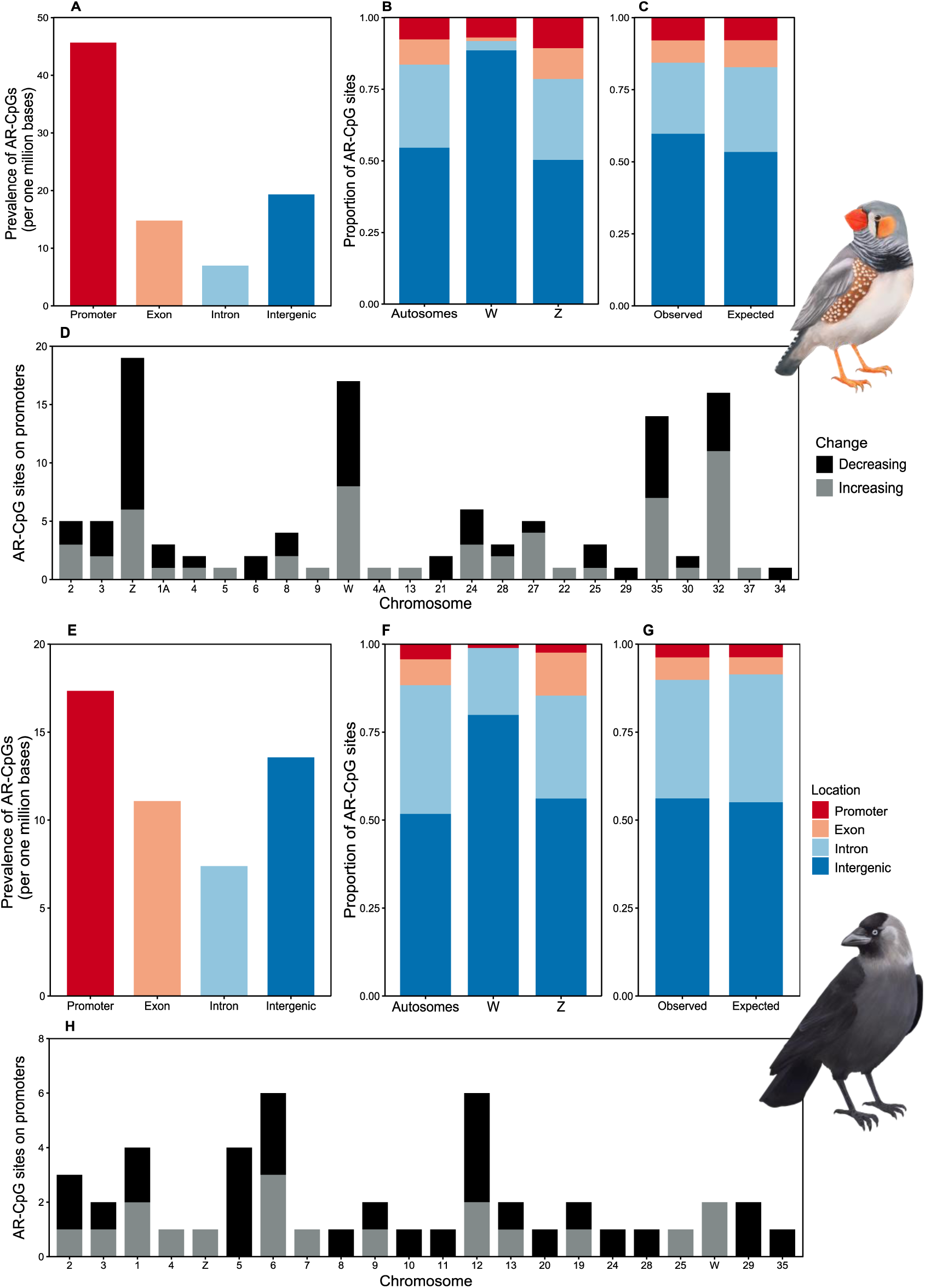
Functional annotation of AR-CpG sites. Count of identified AR-CpG sites per one million base pairs in each annotation category (**A & E**), raw proportion (not normalized) of identified AR-CpG sites in each annotation category by chromosome type (**B & F**) and observed vs. expected in the whole genome (**C & G**) for the zebra finch and jackdaw respectively. Count of AR-CpG sites increasing or decreasing in the degree of DNAm in promoters per chromosome (ordered by decreasing length) for the zebra finch and jackdaw, respectively **(D & H**).

Significant enrichment for AR-CpG sites was observed in both species across annotation categories and chromosome types (Fig.5, Table S5). In the zebra finch, AR-CpG sites were enriched in promoter regions and depleted in introns across all chromosome types (Fig.5A, Table S5A) while in the jackdaw AR-CpG sites were depleted in introns for all chromosome types (Fig.5B, Table S5B). Importantly, in both species, the W chromosome showed the strongest divergence and AR-CpG sites were enriched in intergenic regions, with the highest observed odds ratios (Fig.5, Table S5).

**Figure 5.**
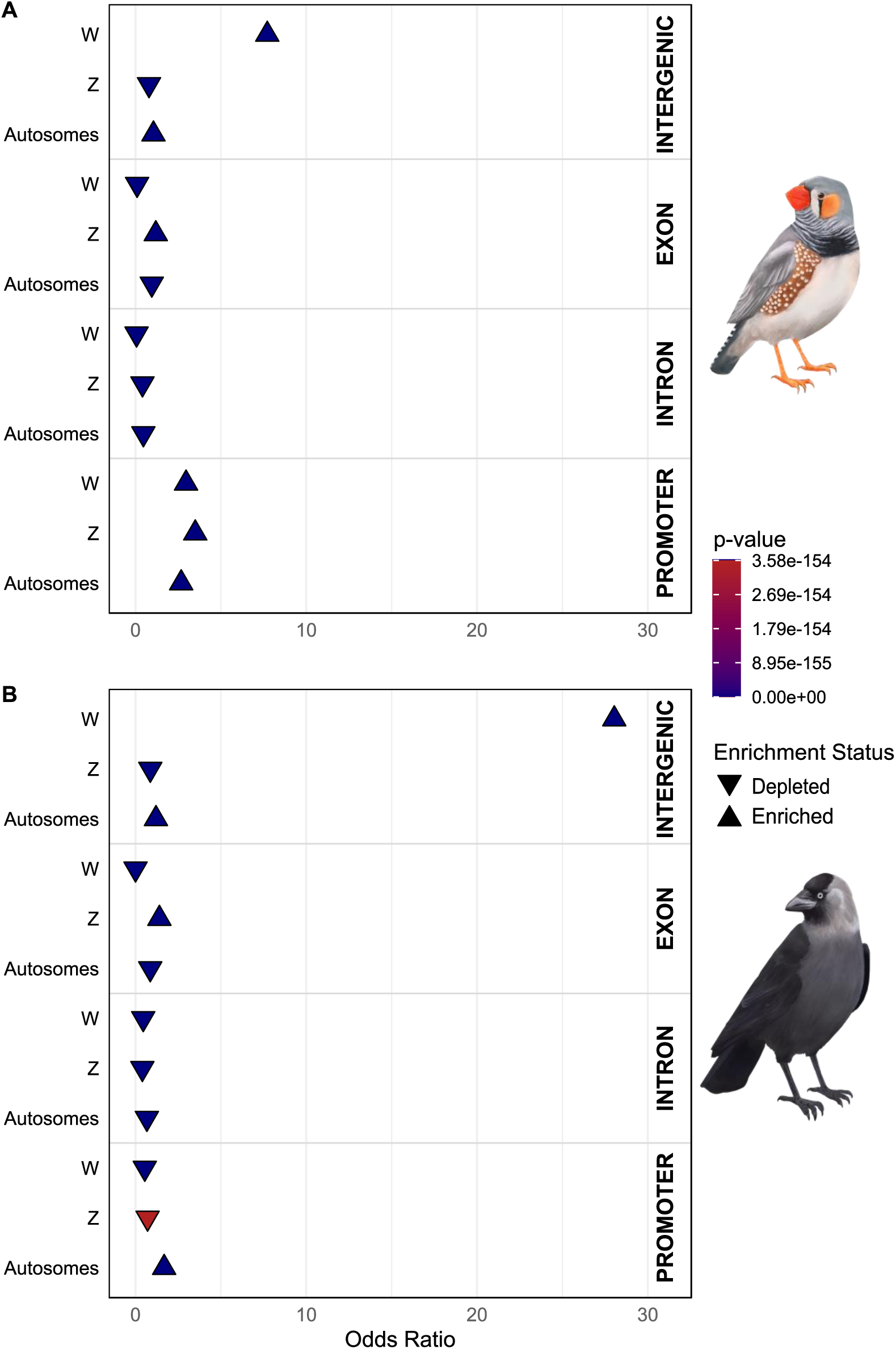
Results of the enrichment analysis across genomic locations and chromosome types in the zebra finch (**A**) and the jackdaw (**B**). Shapes represent the enrichment status of each category, while the color indicates the associated p-value of the Fisher’s exact test.

DNAm within promoter regions is known to block gene expression (Moore et al., 2013; Steyaert et al., 2016), driving phenotypic variation in development, aging, and disease and making it particularly relevant for aging research and a wide range of other biological studies. On the zebra finch sex chromosome promoters, DNAm increased with age on 12 sites and decreased on 21 sites (Fig. 4D). In the jackdaw, all AR-CpG sites in promoters of sex chromosomes increased with age (n=4; Fig. 4H). In the zebra finch autosomal chromosomes, 80 AR-CpG sites were detected in promoter regions, with DNAm increasing with age at 44 sites, and decreasing with age at 36 sites (Fig. 4D). In the jackdaw autosomal chromosomes, 43 AR-CpG sites were found in promoters among which 15 and 28 increased and decreased in DNAm with age, respectively (Fig. 4H). For a small number of AR-CpG sites in promoters the specific genes could be identified (Table S6). In both species, DNAm decreased with age on the majority of intergenic W chromosome AR-CpG sites (Fig.S1). Details of the direction of change in the degree of DNAm at AR-CpG sites per annotation category per chromosome type in each species are depicted in Fig. S1.

The GO enrichment analysis of the genes whose promoters contained AR-CpG sites were not involved in shared biological processes or molecular functions for either species.

The number of AR-CpG sites identified by our pipeline depends on the stringency of the selection criteria, raising the question whether this biases our findings. We therefore repeated the analysis with less stringent criteria, resulting in an increase in identified AR-CpG sites (Fig. S2, S3). However, the overall trends regarding the distribution of AR-CpG sites across the chromosomes remained the same in both species (Fig. S2, S3), leading us to assume that our results were robust with respect to the specific choice of pipeline criteria.

## Discussion

Our finding of a high density of AR-CpG sites on the W chromosome in two avian species highlights the potential of sex chromosome DNAm to drive sex-specific aging (G. Li et al., 2022; Lund et al., 2020). A unique strength of our study was the longitudinal character of the data, combined with whole-genome methylome data, which provided us with a high-resolution view of the entire methylome. Consequently, our whole-genome, longitudinal approach enabled us to detect previously unidentified sex-chromosome-specific DNAm patterns, unlike earlier attempts that were restricted to targeting specific genomic regions (Meyer et al., 2023) or relied on cross-sectional analyses (Sun et al., 2021). We hypothesize this pattern may contribute to sex-dependent aging (e.g. Briga et al., 2017). The similarity of our findings between jackdaws and zebra finches, such as the comparable overall number of AR-CpG sites, their increase and decrease in DNAm over time as well as their distribution within each annotation category on the W chromosome suggest our findings may apply generally to passerine birds.

Whether DNAm increases (hypermethylation) or decreases (hypomethylation) with age typically varies among CpG sites (S. Li et al., 2017). On the W chromosome in both species, DNAm decreased with age in the majority of AR-CpG sites. The avian W chromosome contains a high density of transposable elements (Peona et al., 2021), whose expression has negative fitness consequences (Warmuth et al., 2022), which may explain why DNAm is substantially higher on W compared to the autosomal chromosomes (Tone et al., 1984, also in our study species, data not shown). Decreasing DNAm with age likely leads to declining suppression of transposable element expression, and thereby increased genomic instability and deregulation of gene expression, both drivers of senescence (Gaudet et al., 2003; López-Gil et al., 2023). On the W chromosomes of both species, AR-CpG sites were significantly enriched in intergenic regions. Besides transposons, intergenic regions also harbor microRNAs (Ambros 2004) that modulate longevity and the aging processes (Kinser & Pincus, 2020) while playing a significant role in cell proliferation, cell death, and hematopoiesis (Ambros, 2004). Dysregulation of microRNA gene expression can lead to malignancy (Costinean et al., 2006) and disrupted immune cell homeostasis (Rose et al., 2021). Decreased levels of DNAm with age in intergenic regions of the W chromosome suggests loss of the ability to inhibit detrimental genetic elements such as deleterious mutations leading to genomic instability, which is one of the hallmarks of aging. Our findings suggest that age-related changes in DNAm on the W chromosome may correlate with the shorter lifespans observed in female birds compared to males (Tower & Arbeitman, 2009). However, further longitudinal studies are needed to disentangle developmental and aging-related epigenetic dynamics in this context.

The density of AR-CpG sites was higher than expected by chance on the zebra finch Z chromosome, whereas there were slightly fewer AR-CpG sites than expected on the jackdaw Z chromosome. This difference matches the sex difference in lifespan in the two species: in zebra finches, females have a shorter lifespan (Briga et al., 2017), whereas there is no difference in lifespan between the sexes in jackdaws (Boonekamp et al., 2014). The density of AR-CpG sites deviated significantly from the null expectations on most autosomes, but most deviations were small. The notable exceptions were chromosomes 32 and 35 in the zebra finch, where the density of AR-CpG sites was higher than expected. Whether variation in the AR-CpG density among chromosomes (other than the W chromosome) reflects the frequency of transposable elements or other functional components among chromosomes remains to be investigated.

Our finding that AR-CpG sites on the W chromosome mostly decreased in DNAm with age and that most such sites were located in intergenic regions contrasts research on the human haploid, heterogametic Y chromosome where the majority of AR-CpG sites were found to become hypermethylated with age and were located in genes (Lund et al., 2020). On the other hand, a single early study found the mammalian X chromosome to harbor relatively few AR-CpG sites (Acevedo et al., 2015), as we found for the jackdaw Z chromosome. The avian sex chromosomes ZW are not homologous to the mammalian XY (Nada et al., 1999) and it is therefore unsurprising that DNAm patterns differ from male heterogametic (XY) species. Our findings suggest that DNAm on sex chromosomes likely plays different roles in male and female heterogametic systems. Furthermore, our results underscore the notion that some of the key mechanisms driving sex-specific differences in aging might be fundamental and thus conserved among closely related species, but not across taxa. Consequently, efforts should be directed towards deciphering the role of sex chromosomes in aging across a broader range of species as well as understanding the mechanisms underlying this process in species with various sex determination systems.

## Conclusions

Utilizing longitudinal, whole-genome DNAm data from two avian species, we were able to characterize the distribution of CpG sites whose DNAm significantly changes with age. The location, direction of change in DNAm and functional significance of those lead us to believe that avian sex chromosomes-particularly the haploid, female-specific W- are subject to distinct age-related epigenetic regulation. Furthermore, our findings suggest that the observed significant changes in DNAm can lead to genomic instability, potentially contributing to the observed shorter lifespans of female birds compared to males. While these findings are a first step towards deciphering the role that epigenetic changes of sex chromosomes play in aging, further research across diverse taxa will be essential to uncover conserved mechanisms underlying the aging process and sex-dependent aging.

## Acknowledgments

We thank the Center for Information Technology at the University of Groningen for support and access to the Peregrine and Hábrók high-performance computing clusters. We express our gratitude to Ellis Mulder for her help in the laboratory, Berber Maarsingh for illustrations, and Raphaël Scherrer for assistance with the statistical analyses. We are also indebted to the animal caretakers at the University of Groningen as well as numerous students whose invaluable help made this project possible.

## Ethics approval

All methods and experiments detailed in this manuscript were performed under the approval of the Central Committee for Animal Experiments (Centrale Commissie Dierproeven) of the Netherlands, under licenses AVD1050020174344 and AVD1050020184967.

## Data Accessibility and Benefit-Sharing

The dataset(s) supporting the conclusions of this article is (are) available in the the National Center for Biotechnology Information under BioProject ID PRJNA1108628. The bioinformatics code and R code, as well as intermediate and final data files used in this study can be found at https://doi.org/10.5061/dryad.wm37pvmw8.

## Author contributions

MT, JS, and SV conceived and designed the study. MT and JS performed the bioinformatics and data analysis under the supervision of SV and PJP. FF performed the genomic annotation. MT and SV wrote the manuscript with input from all authors. All authors approved the final version of the manuscript

## Supplementary Material

**Figure S1.**
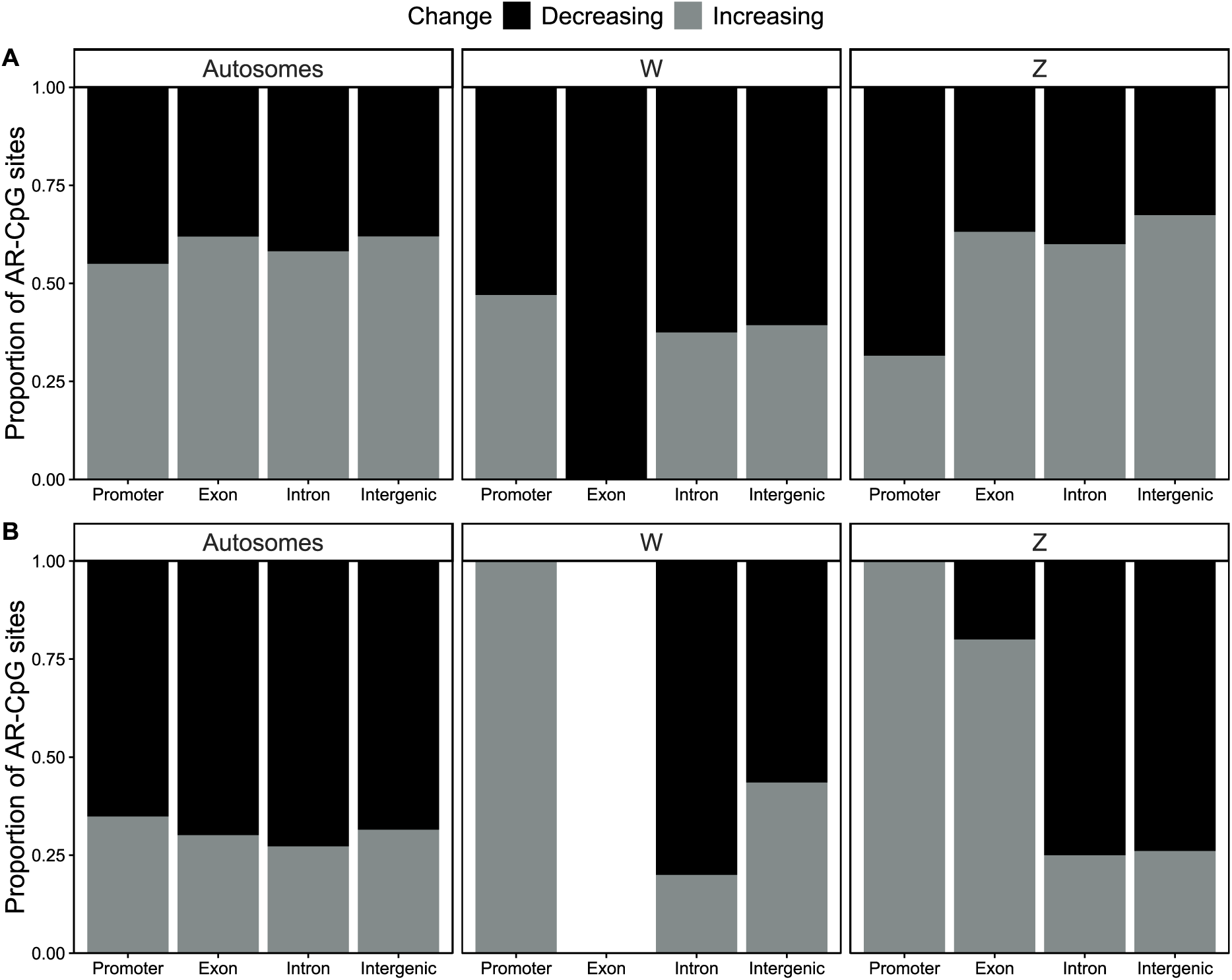
Proportion of AR-CpG sites increasing and decreasing in DNAm with age in each annotation category in the zebra finch (**A**) and the jackdaw (**B**).

**Figure S2.**
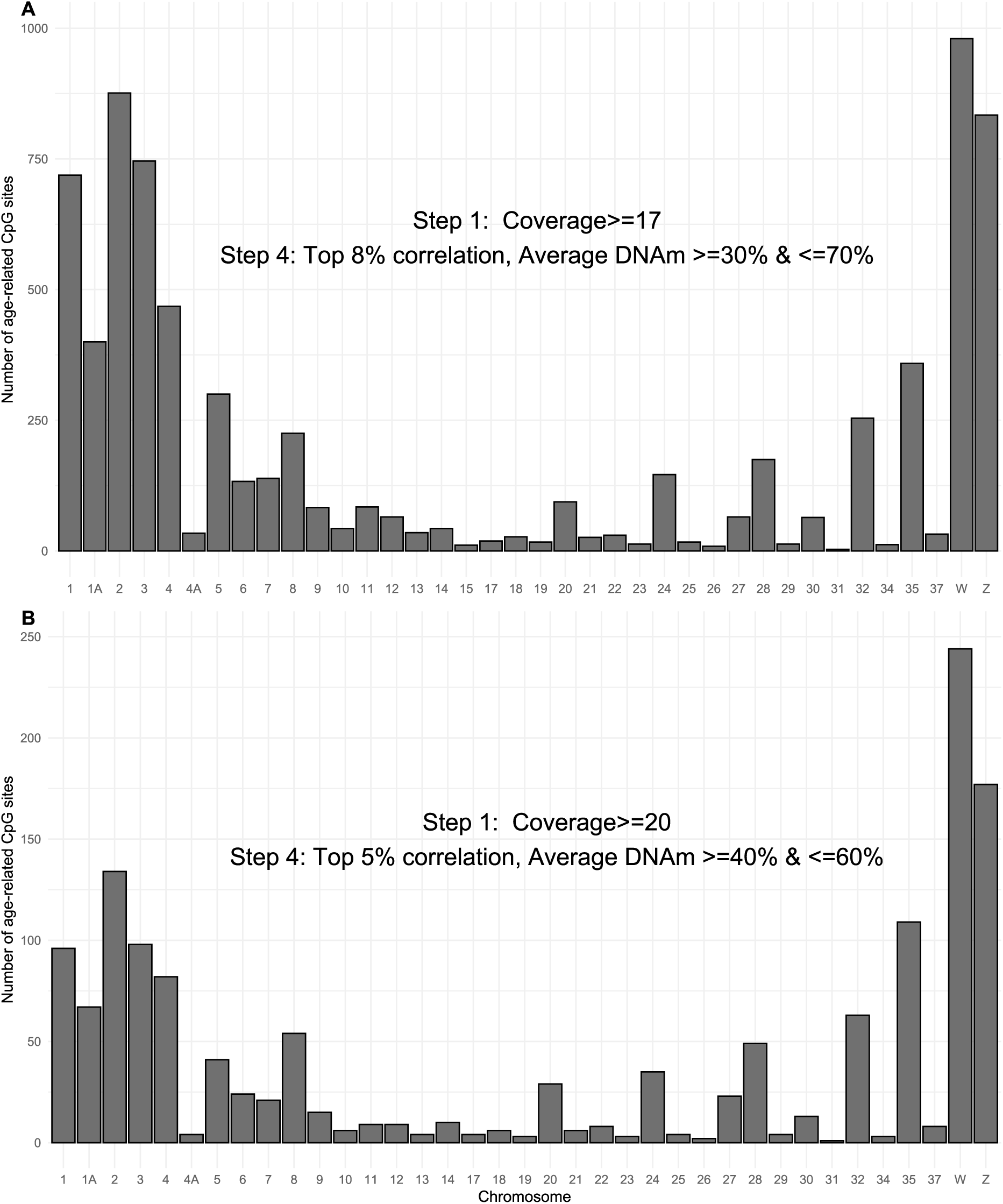
Comparison of “lenient” (A) and “standard” (B) pipelines for the zebra finch. The lenient pipeline selects positions with average coverage of ≥17 (Step 1), sites with average % DNAm between 30% and 70% (Step 2) and at the top 8% of the correlations between % DNAm and Δage per 20 coverage (Step 4). As a result the total number of sites finally selected increases to 7,593 in the zebra finch. Note that the numbers of sites per chromosome are highly correlated between the two pipelines (r_(35)_ =0.95, p<0.001) pointing to the high repeatability and robustness of the pipeline in response to threshold modifications making it applicable to the whole ranges of methylation data.

**Figure S3.**
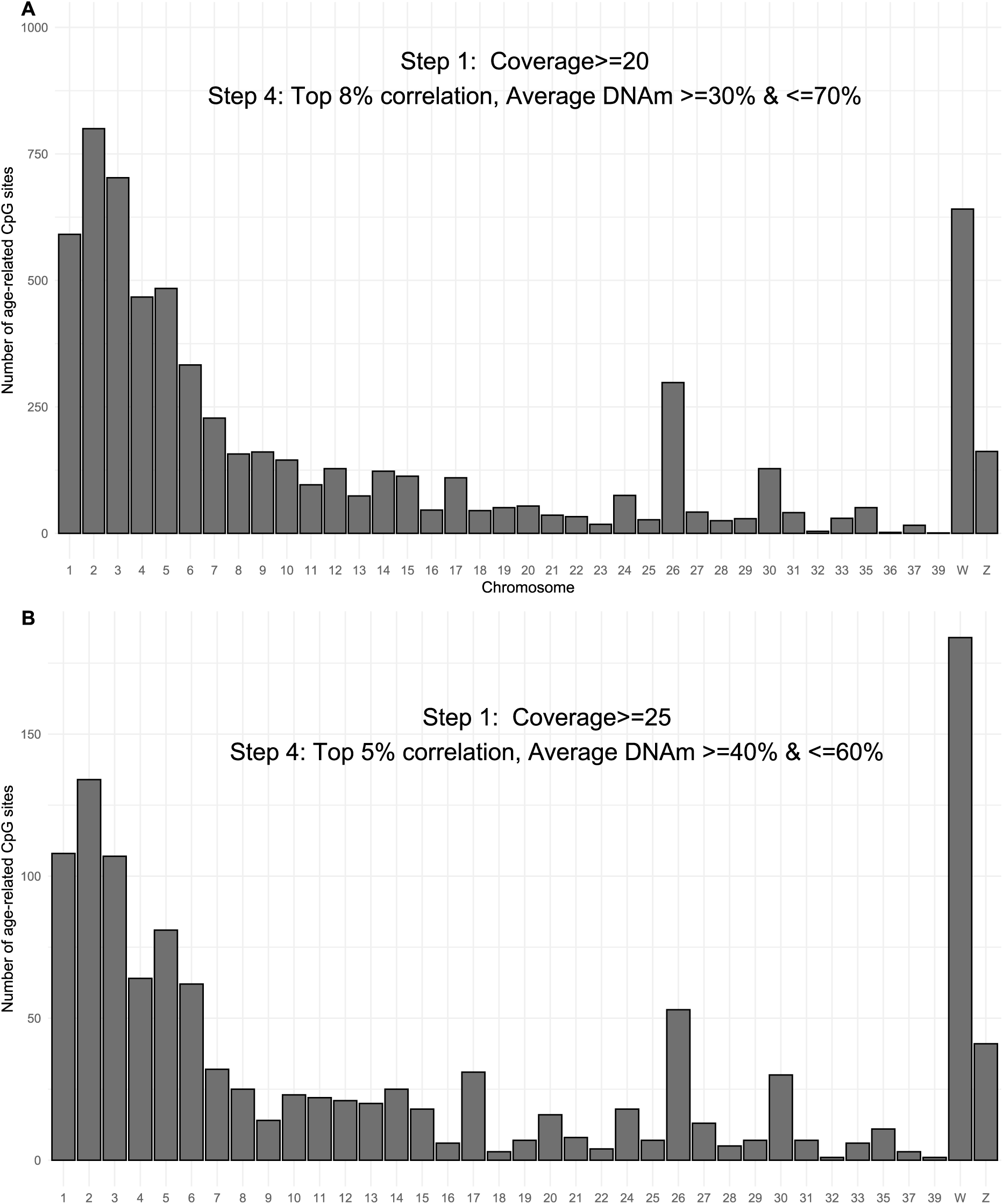
Comparison of “lenient” (A) and “standard” (B) pipelines for the jackdaw. The lenient pipeline selects positions with average coverage of≥20 (Step 1), sites with average % DNAm between 30% and 70% (Step 2) and at the top 8% of the correlations between % DNAm and Δage per 20 coverage (Step 4). As a result the total number of sites finally selected increases to 9, in the zebra finch. Note that the numbers of sites per chromosome are highly correlated between the two pipelines (r_(35)_ =0.95, p<0.001) pointing to the high repeatability and robustness of the pipeline in response to threshold modifications making it applicable to the whole ranges of methylation data.

**Table S1.**
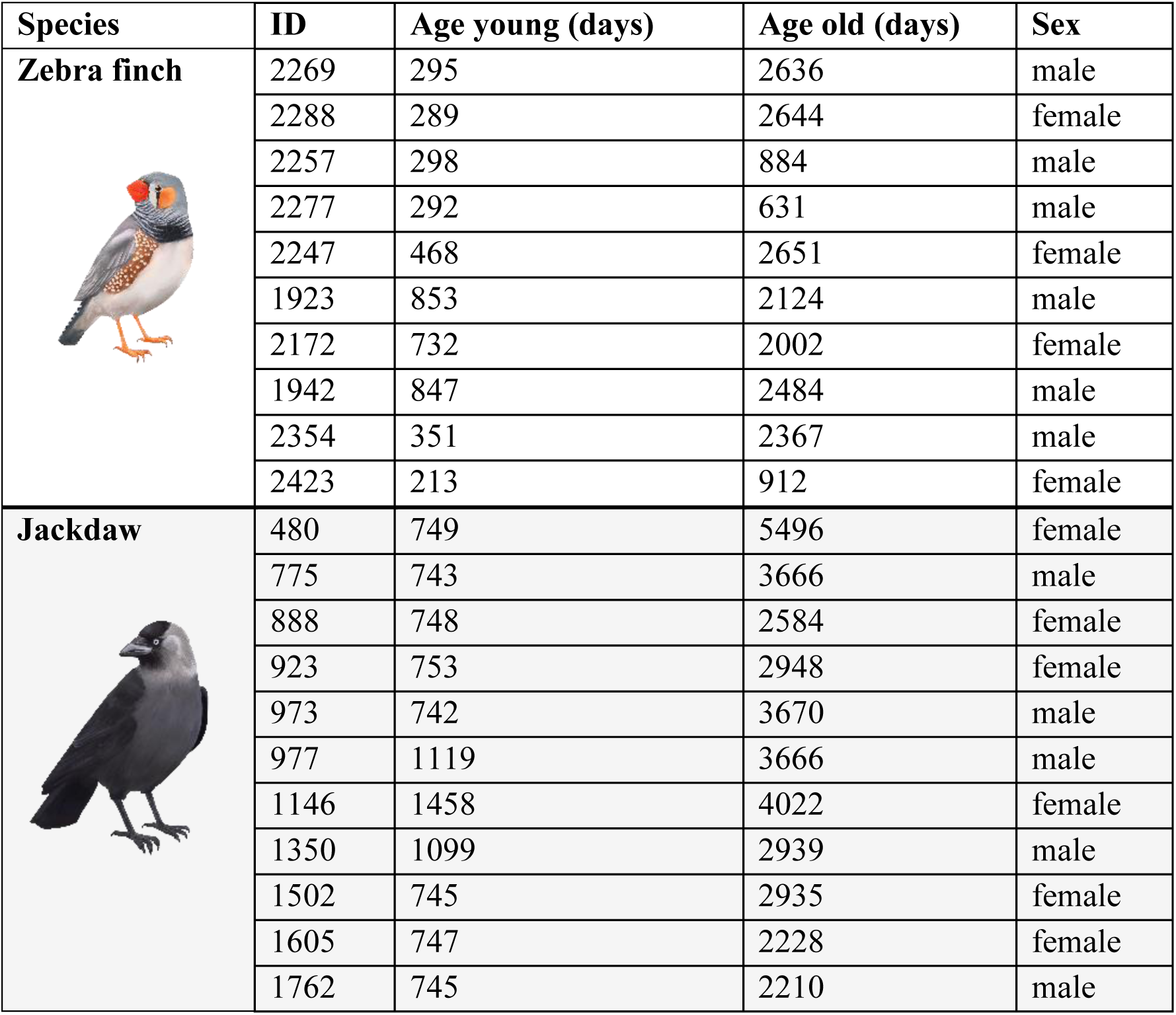
Chronological age at collection and sex of all the individuals in our dataset.

| Species | ID | Age young (days) | Age old (days) | Sex |
| --- | --- | --- | --- | --- |
| **Zebra finch** | 2269 | 295 | 2636 | male |
|  | 2288 | 289 | 2644 | female |
|  | 2257 | 298 | 884 | male |
|  | 2277 | 292 | 631 | male |
|  | 2247 | 468 | 2651 | female |
|  | 1923 | 853 | 2124 | male |
|  | 2172 | 732 | 2002 | female |
|  | 1942 | 847 | 2484 | male |
|  | 2354 | 351 | 2367 | male |
|  | 2423 | 213 | 912 | female |
| **Jackdaw** | 480 | 749 | 5496 | female |
|  | 775 | 743 | 3666 | male |
|  | 888 | 748 | 2584 | female |
|  | 923 | 753 | 2948 | female |
|  | 973 | 742 | 3670 | male |
|  | 977 | 1119 | 3666 | male |
|  | 1146 | 1458 | 4022 | female |
|  | 1350 | 1099 | 2939 | male |
|  | 1502 | 745 | 2935 | female |
|  | 1605 | 747 | 2228 | female |
|  | 1762 | 745 | 2210 | male |

**Table S2.**
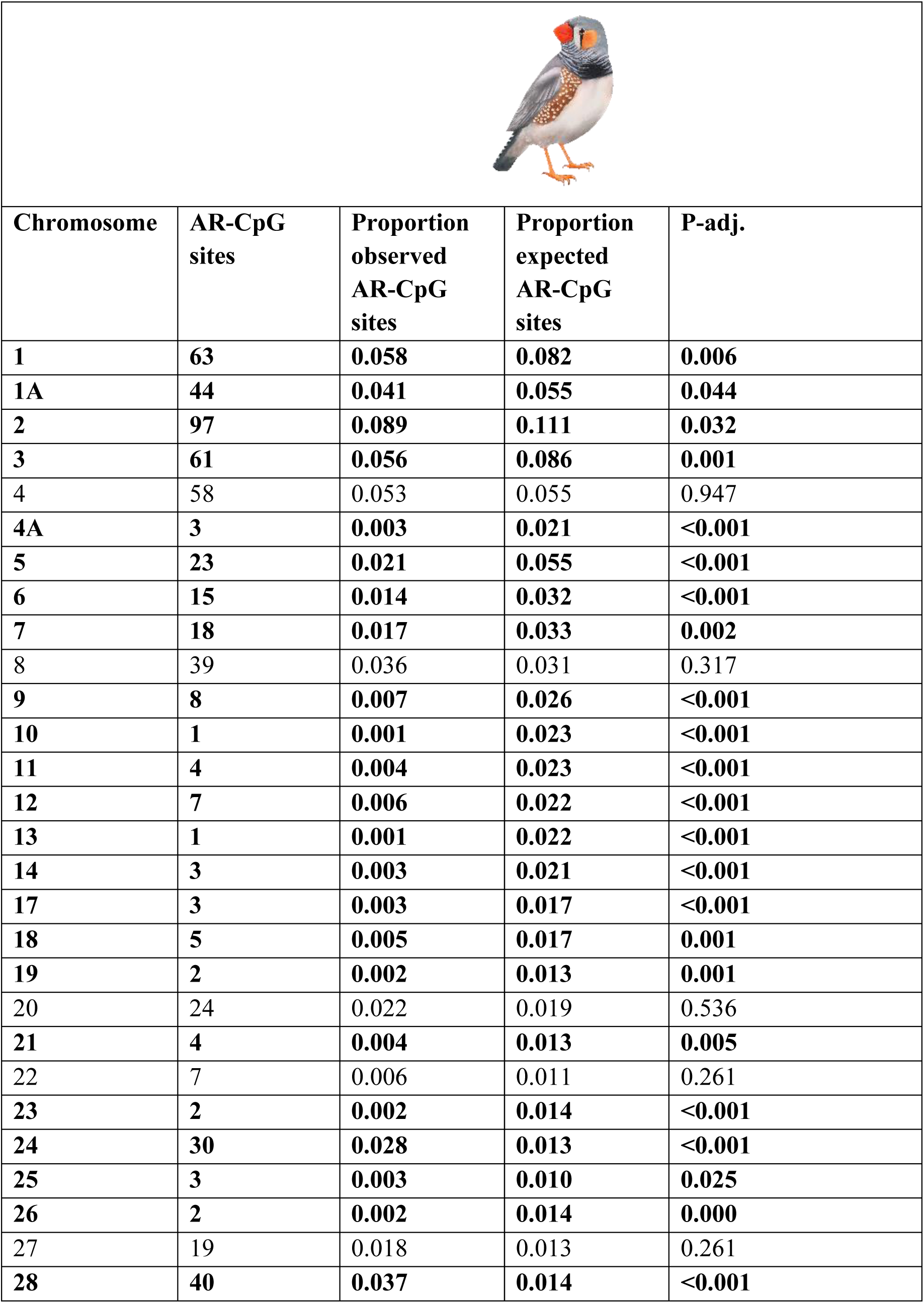
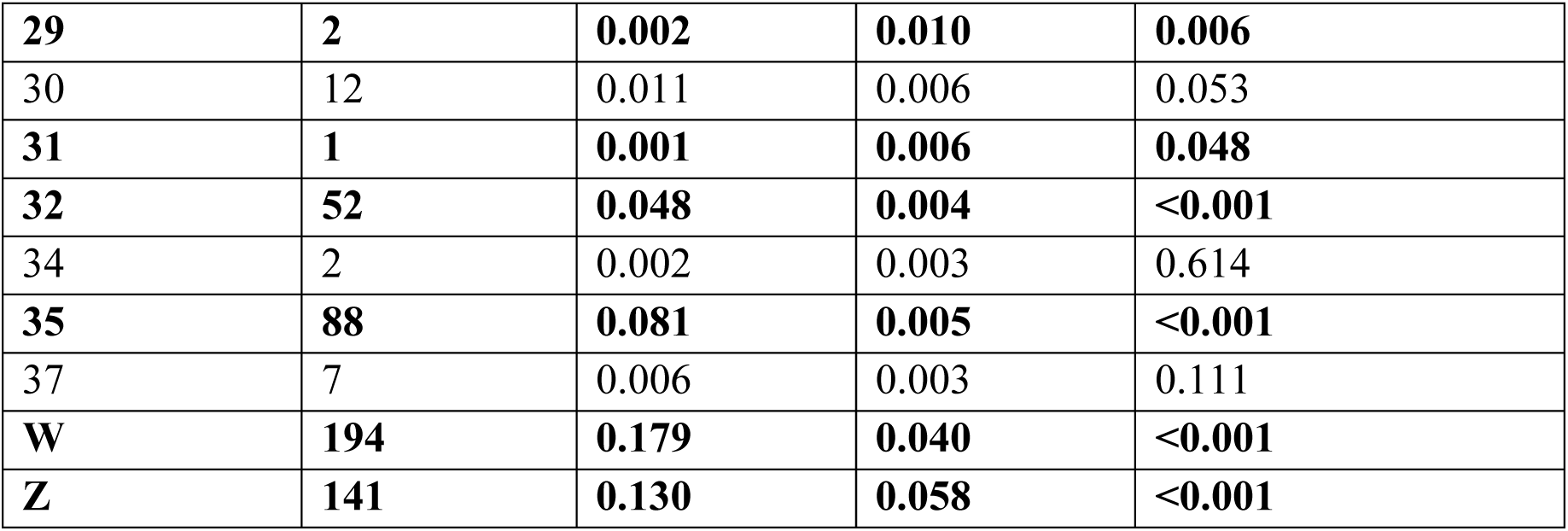
Proportions of observed and expected distribution of age-related CpG sites in each chromosome and results of two-sided exact binomial tests of observed vs. expected proportions of AR-CpG sites in each chromosome in the zebra finch. Significant differences (p<0.05) marked in bold.

|  |  |  |  |  |
| --- | --- | --- | --- | --- |
| Chromosome | AR-CpG sites | Proportion observed AR-CpG sites | Proportion expected AR-CpG sites | P-adj. |
| **1** | **63** | **0.058** | **0.082** | **0.006** |
| **1A** | **44** | **0.041** | **0.055** | **0.044** |
| **2** | **97** | **0.089** | **0.111** | **0.032** |
| **3** | **61** | **0.056** | **0.086** | **0.001** |
| 4 | 58 | 0.053 | 0.055 | 0.947 |
| **4A** | **3** | **0.003** | **0.021** | **<0.001** |
| **5** | **23** | **0.021** | **0.055** | **<0.001** |
| **6** | **15** | **0.014** | **0.032** | **<0.001** |
| **7** | **18** | **0.017** | **0.033** | **0.002** |
| 8 | 39 | 0.036 | 0.031 | 0.317 |
| **9** | **8** | **0.007** | **0.026** | **<0.001** |
| **10** | **1** | **0.001** | **0.023** | **<0.001** |
| **11** | **4** | **0.004** | **0.023** | **<0.001** |
| **12** | **7** | **0.006** | **0.022** | **<0.001** |
| **13** | **1** | **0.001** | **0.022** | **<0.001** |
| **14** | **3** | **0.003** | **0.021** | **<0.001** |
| **17** | **3** | **0.003** | **0.017** | **<0.001** |
| **18** | **5** | **0.005** | **0.017** | **0.001** |
| **19** | **2** | **0.002** | **0.013** | **0.001** |
| 20 | 24 | 0.022 | 0.019 | 0.536 |
| **21** | **4** | **0.004** | **0.013** | **0.005** |
| 22 | 7 | 0.006 | 0.011 | 0.261 |
| **23** | **2** | **0.002** | **0.014** | **<0.001** |
| **24** | **30** | **0.028** | **0.013** | **<0.001** |
| **25** | **3** | **0.003** | **0.010** | **0.025** |
| **26** | **2** | **0.002** | **0.014** | **0.000** |
| 27 | 19 | 0.018 | 0.013 | 0.261 |
| **28** | **40** | **0.037** | **0.014** | **<0.001** |
| **29** | **2** | **0.002** | **0.010** | **0.006** |
| 30 | 12 | 0.011 | 0.006 | 0.053 |
| **31** | **1** | **0.001** | **0.006** | **0.048** |
| **32** | **52** | **0.048** | **0.004** | **<0.001** |
| 34 | 2 | 0.002 | 0.003 | 0.614 |
| **35** | **88** | **0.081** | **0.005** | **<0.001** |
| 37 | 7 | 0.006 | 0.003 | 0.111 |
| **W** | **194** | **0.179** | **0.040** | **<0.001** |
| **Z** | **141** | **0.130** | **0.058** | **<0.001** |

**Table S3.**
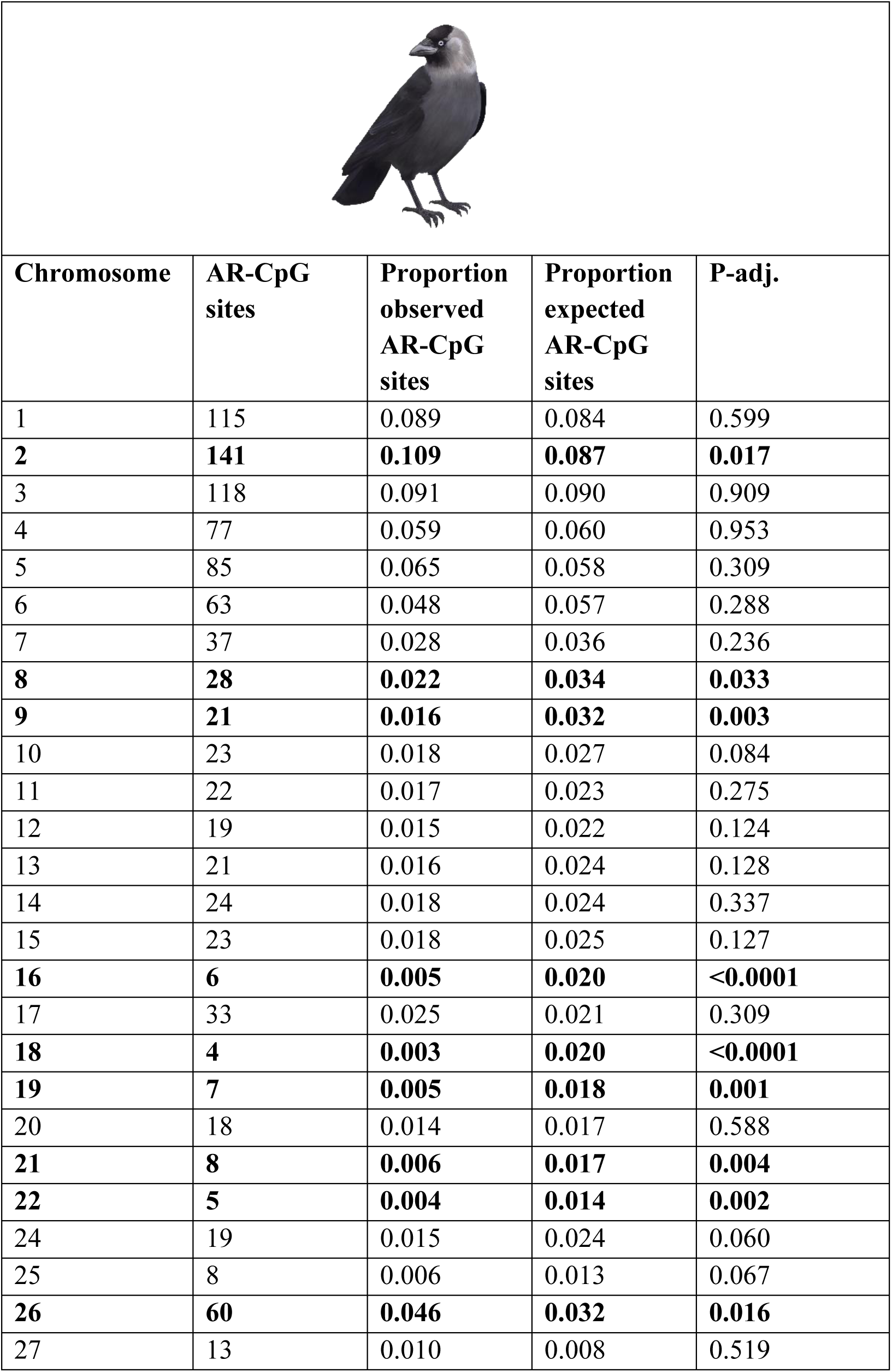
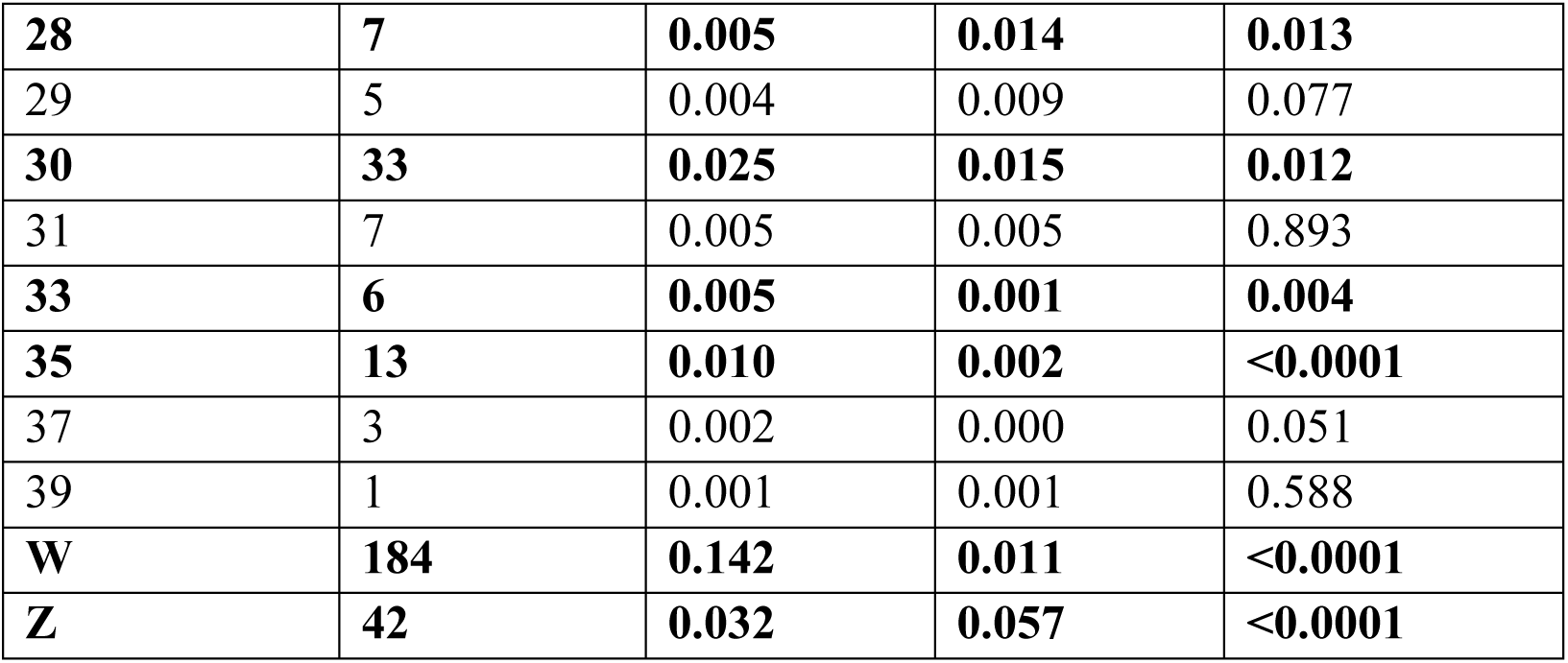
Proportions of observed and expected distribution of age-related CpG sites in each chromosome and results of two-sided exact binomial tests of observed vs. expected proportions of AR-CpG sites in each chromosome in the jackdaw. Significant differences (p<0.05) marked in bold.

|  |  |  |  |  |
| --- | --- | --- | --- | --- |
| Chromosome | AR-CpG sites | Proportion observed AR-CpG sites | Proportion expected AR-CpG sites | P-adj. |
| 1 | 115 | 0.089 | 0.084 | 0.599 |
| **2** | **141** | **0.109** | **0.087** | **0.017** |
| 3 | 118 | 0.091 | 0.090 | 0.909 |
| 4 | 77 | 0.059 | 0.060 | 0.953 |
| 5 | 85 | 0.065 | 0.058 | 0.309 |
| 6 | 63 | 0.048 | 0.057 | 0.288 |
| 7 | 37 | 0.028 | 0.036 | 0.236 |
| **8** | **28** | **0.022** | **0.034** | **0.033** |
| **9** | **21** | **0.016** | **0.032** | **0.003** |
| 10 | 23 | 0.018 | 0.027 | 0.084 |
| 11 | 22 | 0.017 | 0.023 | 0.275 |
| 12 | 19 | 0.015 | 0.022 | 0.124 |
| 13 | 21 | 0.016 | 0.024 | 0.128 |
| 14 | 24 | 0.018 | 0.024 | 0.337 |
| 15 | 23 | 0.018 | 0.025 | 0.127 |
| **16** | **6** | **0.005** | **0.020** | **<0.0001** |
| 17 | 33 | 0.025 | 0.021 | 0.309 |
| **18** | **4** | **0.003** | **0.020** | **<0.0001** |
| **19** | **7** | **0.005** | **0.018** | **0.001** |
| 20 | 18 | 0.014 | 0.017 | 0.588 |
| **21** | **8** | **0.006** | **0.017** | **0.004** |
| **22** | **5** | **0.004** | **0.014** | **0.002** |
| 24 | 19 | 0.015 | 0.024 | 0.060 |
| 25 | 8 | 0.006 | 0.013 | 0.067 |
| **26** | **60** | **0.046** | **0.032** | **0.016** |
| 27 | 13 | 0.010 | 0.008 | 0.519 |
| **28** | **7** | **0.005** | **0.014** | **0.013** |
| 29 | 5 | 0.004 | 0.009 | 0.077 |
| **30** | **33** | **0.025** | **0.015** | **0.012** |
| 31 | 7 | 0.005 | 0.005 | 0.893 |
| **33** | **6** | **0.005** | **0.001** | **0.004** |
| **35** | **13** | **0.010** | **0.002** | **<0.0001** |
| 37 | 3 | 0.002 | 0.000 | 0.051 |
| 39 | 1 | 0.001 | 0.001 | 0.588 |
| **W** | **184** | **0.142** | **0.011** | **<0.0001** |
| **Z** | **42** | **0.032** | **0.057** | **<0.0001** |

**Table S4.** Proportions of observed and expected distribution of age-related CpG sites in each annotation category and results of two-sided exact binomial tests of observed vs. expected proportions of AR-CpG sites in each annotation category for the zebra finch (**A**) and the jackdaw (**B**). Expected proportions were calculated from the location of all CpG sites captured by our analysis in each annotation category.

| *Category* | *A* |  |  | *B* |  |  |
| --- | --- | --- | --- | --- | --- | --- |
|  | *Observed proportions* | *Expected proportions* | *p-value (two-sided)* | *Observed proportions* | *Expected proportions* | *p-value (two-sided)* |
| *Promoters* | 0.079 | 0.094 | **0.044** | 0.038 | 0.037 | **0.02** |
| *Exons* | 0.078 | 0.534 | **<0.001** | 0.064 | 0.049 | 0.471 |
| *Introns* | 0.247 | 0.294 | **<0.001** | 0.337 | 0.363 | 0.06 |
| *Intergenic regions* | 0.598 | 0.078 | **<0.001** | 0.562 | 0.551 | 0.879 |

**Table S5.**
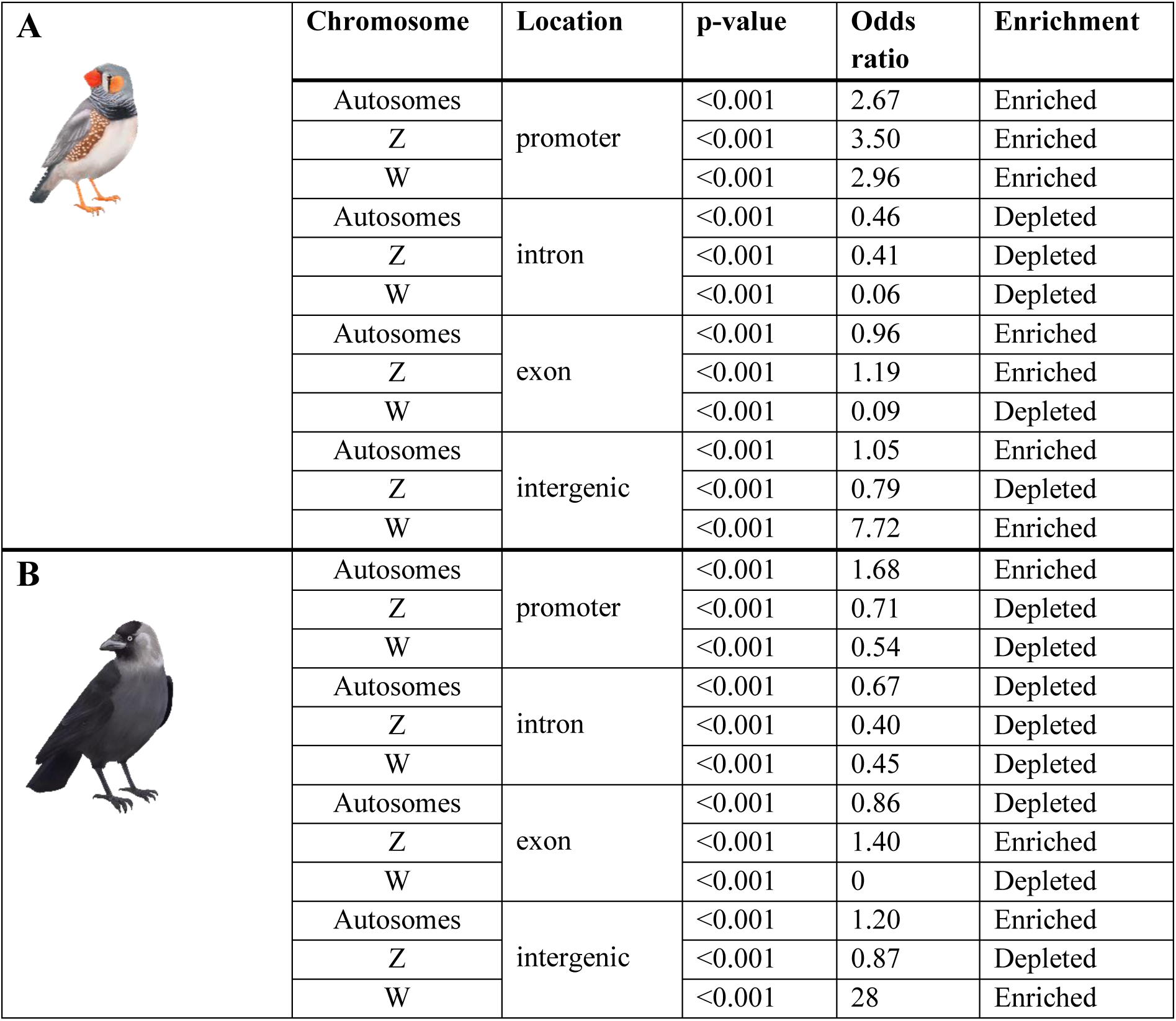
Enrichment and depletion of AR-CpG sites across genomic relative to total genomic coverage in the autosomes grouped together, the W and Z chromosomes of the zebra finch (**A**) and jackdaw (**B**). Enrichment statistics were calculated using a Fisher’s exact test. Here enrichment is defined as an odds ratio >1 and depletion as <1.

| **A** | Chromosome | Location | p-value | Odds ratio | Enrichment |
| --- | --- | --- | --- | --- | --- |
|  | Autosomes | promoter | <0.001 | 2.67 | Enriched |
|  | Z |  | <0.001 | 3.50 | Enriched |
|  | W |  | <0.001 | 2.96 | Enriched |
|  | Autosomes | intron | <0.001 | 0.46 | Depleted |
|  | Z |  | <0.001 | 0.41 | Depleted |
|  | W |  | <0.001 | 0.06 | Depleted |
|  | Autosomes | exon | <0.001 | 0.96 | Enriched |
|  | Z |  | <0.001 | 1.19 | Enriched |
|  | W |  | <0.001 | 0.09 | Depleted |
|  | Autosomes | intergenic | <0.001 | 1.05 | Enriched |
|  | Z |  | <0.001 | 0.79 | Depleted |
|  | W |  | <0.001 | 7.72 | Enriched |
| **B** | Autosomes | promoter | <0.001 | 1.68 | Enriched |
|  | Z |  | <0.001 | 0.71 | Depleted |
|  | W |  | <0.001 | 0.54 | Depleted |
|  | Autosomes | intron | <0.001 | 0.67 | Depleted |
|  | Z |  | <0.001 | 0.40 | Depleted |
|  | W |  | <0.001 | 0.45 | Depleted |
|  | Autosomes | exon | <0.001 | 0.86 | Depleted |
|  | Z |  | <0.001 | 1.40 | Enriched |
|  | W |  | <0.001 | 0 | Depleted |
|  | Autosomes | intergenic | <0.001 | 1.20 | Enriched |
|  | Z |  | <0.001 | 0.87 | Depleted |
|  | W |  | <0.001 | 28 | Enriched |

**Table S6.**
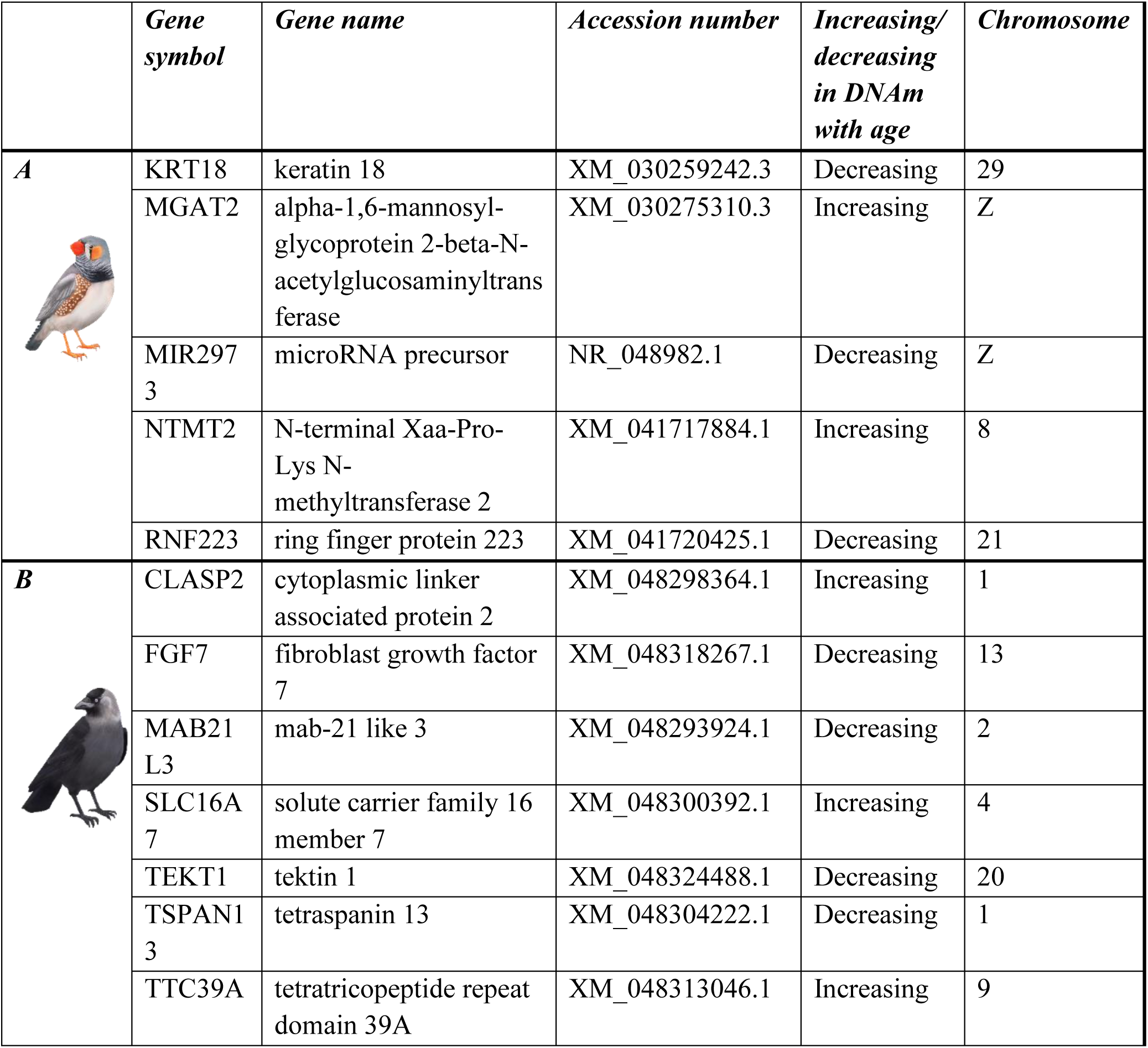
Identified genes in whose promoters we located AR-CpG sites and whether DNAm increased or decreased with time at that AR-CpG site in the zebra finch (**A**) and jackdaw (**B**). The 88 and 43 remaining AR-CpG sites in promoters for the zebra finch and jackdaw respectively were located in genes with unknown function.

|  | *Gene symbol* | *Gene name* | *Accession number* | *Increasing/*<br>*decreasing*<br>*in DNAm*<br>*with age* | *Chromosome* |
| --- | --- | --- | --- | --- | --- |
| **A** | KRT18 | keratin 18 | XM_030259242.3 | Decreasing | 29 |
|  | MGAT2 | alpha-1,6-mannosyl-glycoprotein 2-beta-N-acetylglucosaminyltransferase | XM_030275310.3 | Increasing | Z |
|  | MIR2973 | microRNA precursor | NR_048982.1 | Decreasing | Z |
|  | NTMT2 | N-terminal Xaa-Pro-Lys N-methyltransferase 2 | XM_041717884.1 | Increasing | 8 |
|  | RNF223 | ring finger protein 223 | XM_041720425.1 | Decreasing | 21 |
| **B** | CLASP2 | cytoplasmic linker associated protein 2 | XM_048298364.1 | Increasing | 1 |
|  | FGF7 | fibroblast growth factor 7 | XM_048318267.1 | Decreasing | 13 |
|  | MAB21L3 | mab-21 like 3 | XM_048293924.1 | Decreasing | 2 |
|  | SLC16A7 | solute carrier family 16 member 7 | XM_048300392.1 | Increasing | 4 |
|  | TEKT1 | tektin 1 | XM_048324488.1 | Decreasing | 20 |
|  | TSPAN13 | tetraspanin 13 | XM_048304222.1 | Decreasing | 1 |
|  | TTC39A | tetratricopeptide repeat domain 39A | XM_048313046.1 | Increasing | 9 |

